# Exploring vitamin D and its analogues as a possible therapeutic intervention in targeting inflammatory mediators in neurodegenerative disorders like Parkinson’s disease

**DOI:** 10.1101/2025.01.10.632401

**Authors:** Randrita Pal, Nilansu Das, Barnali Ray Basu

**Author notes:** Corresponding author: **Barnali Ray Basu, An active e-mail address of the corresponding author:**.

## Abstract

Parkinson’s disease characterized by dopaminergic neuronal death has been linked with neuroinflammatory processes which involve self-perpetuating detrimental events leading to prolonged neurodegeneration. Vitamin D has been found to possess neuroprotective and immuno-modulatory effects which might justify elevated inflammation and hypovitaminosis D in PD patients. So, we aimed to find the possible interactions of Vitamin-D and its analogues with inflammatory mediators prevalent in PD to get more insight into Vitamin-D’s role in neuroinflammation by assessing specific interactions between Vitamin-D and its marketed analogues with different inflammatory mediators. Moreover, revealing a possible drug repurposing with Vitamin-D analogues against neuroinflammation and other inflammatory conditions.

Molecular docking was conducted with 10 inflammatory markers (p38 MAPK, NF-κB, iNOS, nNOS, COX-2, TNFα, IL6, TNFα, IL1β, CRP) with bio-active form of Vitamin-D (calcitriol) and its analogues. Among the studied ligands, calcitriol, calcipotriol, seocalcitol, paricalcitol and falecalcitriol were found to be potential inhibitors of different inflammatory agents. Specific standard drugs or experimental inhibitors were used to compared the binding affinity between protein and ligand. BBB permeability of the tested ligands also checked as well as neurotoxicity. Most of the ligands were found to be BBB permeable possibly interacting with inflammatory mediators within the brain and were also found to be non-neurotoxic.

This study sheds light on VD and its analogues being possible candidates against various inflammatory markers. The findings could further be used as stepping stone for new study on repurposing of these analogues in neuroinflammation and other inflammatory disease conditions.

## 1. Introduction

Parkinson’s disease (PD) onset is orchestrated through abnormal accumulation of α-synuclein in the dopaminergic neurons (substantia nigra). Neuroinflammation plays an apex role in PD pathogenesis. The neuroinflammatory response however, is a complex scenario involving microglial activation, astrocytes, gut microbiota and also pathogenic factors which can cross the blood-brain-barrier (BBB) to create pro-inflammatory cytokine inflation (Liu *et al,* 2022).

Neuroinflammation is a resultant of malfunction of various associated proteins. The p38 Mitogen-Activated Protein Kinase (MAPK) pathway and Nuclear Factor-kappa B (NF-κB) are two critical signalling pathways involved in neuroinflammation. Both pathways play significant roles in the brain’s inflammatory response, contributing to various neurological disorders, including Parkinson’s disease (Shih *et al*, 2015). These two key molecules contribute to neuroinflammation by three-way mechanism. Both commence microglial activation by modulating expression of pro-inflammatory cytokines, TNF-α, IL-6, IL-1β, and chemokines (Lau *et al*, 2021). Apart from pro-inflammatory cytokines, NF-κB is also known to produce enzymes like cyclooxygenase-2 (COX-2) (Shih *et al*, 2015). MAPK is also reported to stimulate neurotoxic mediators like nitric oxides (NO), which further aggravate neuronal damage. The third hit is that it triggers apoptosis and cause instability of the BBB. p38 and NF-κB are reported to co-activate each other with activated NF-κB upregulating cytokine production and further activating p38 creating a positive feedback loop (Zhao *et al*, 2019).

Inflammatory stimuli like cytokines induce production of Inducible Nitric Oxide Synthase (iNOS) in microglia, astrocytes, and neurons. iNOS produces NO inducing oxidative stress and neuronal damage causing neuroinflammation. Additionally, NO reacts with superoxide to form peroxynitrite. Neuronal Nitric Oxide Synthase (nNOS) is typically constitutively expressed in neurons and plays a role in normal neuronal functions such as synaptic plasticity, neurotransmission, and vascular regulation. While low levels of NO provide neuroprotection, promoting synaptic plasticity and increased blood flow, inflammatory responses produce high levels of NO which can exacerbate neuroinflammation and neuronal injury. This is particularly problematic when nNOS interacts with iNOS-produced NO, leading to peroxynitrite formation (Picón-Pagès *et al*, 2019), a highly reactive species that can damage lipids, proteins, and DNA, leading to neuronal injury and death.

Cyclo-oxygenase 2 (COX-2), usually present in the body under normal conditions is responsible for formation of prostaglandins from arachidonic acid. However, under stress and cytokine reflexes, their levels increase. During inflammation, COX-2 is elevated in microglia and astrocytes which cause increased prostaglandins, particularly PGE2 which further amplifies neuroinflammatory responses including vasodilation and pain leading to chronic inflammation (Bartels and Leenders, 2010).

TNF-α, IL-6, IL-1β are pro-inflammatory cytokines which initiate inflammation in microglia, astrocytes and even neurons, while CRP is an acute-phase protein produced by glial cells and liver. TNF-α is particularly known to disrupt the BBB, induce oxidative stress and apoptosis. IL-6 is known to have both pro-inflammatory and anti-inflammatory properties, usually activated in presence of other pro-inflammatory cytokines like TNF-α and IL-1β (Al-Roub *et al*, 2021). IL-6 is a key mediator of the acute phase response in the brain as a result of which CRP is produced for immune-modulation. Although IL-6 is useful in infections or injuries, overproduction may lead to exacerbated neuroinflammation, ultimately leading to neuronal death.

Calcitriol, the bioactive form of Vitamin D is a naturally occurring neuroprotective agent and reported to ameliorate neurotoxic processes (Cui *et al*, 2021). It is also reported to exhibit several anti-inflammatory effects such as inhibiting p38 kinase signalling, suppressing prostaglandin action, and subsequently producing pro-inflammatory cytokines and lastly inhibition of NF-κB signalling (Krishnan and Feldman, 2011).

Hypovitaminosis D was observed in Parkinson’s disease patients and supplementation was found to be useful. Studies on animal model showed that VD significantly attenuated microglial cell activation (Iba1-immunoreactive), iNOS and TLR-4 expression, typical hallmarks of the pro-inflammatory (M1) activation of microglia. Additionally, VD was able to decrease pro-inflammatory cytokines mRNA expression in distinct brain areas of the studied animal group. Notably, the anti-inflammatory property of VD in the mouse model was shown where it upregulated the mRNA expression of the anti-inflammatory cytokines (IL-10, IL-4 and TGF-β) as well as increasing the expression of CD163, CD206 and CD204, typical hallmarks of alternative activation of microglia for anti-inflammatory signalling (M2) (Calvello *et al*, 2016). Many Vitamin D analogues are available which are used for treating different ailments. Doxercalciferol is reported to be used against cell senescence and skin aging, secondary hyperthyroidism (Ge *et al*, 2024; Dennis and Albertson, 2006). Prodrugs like doxercalciferol and alfacalcidol are used for hypocalcemia, chronic renal failure, hypoparathyroidism and osteoporosis and hyperthyroidism (Kubodera, 2009). Alfacalcidol, calcitriol, doxercalciferol, falecalcitriol, maxacalcitol and paricalcitol were used as Vitamin D Receptor activators (Zhang *et al*, 2016) and were seen effective against chronic kidney disease (Palmer *et al*, 2009). Seocalcitol has been reported to have anti-cancer use (Beuno *et al*,2022) and also against certain kidney ailments.

Here, our primary target was not only to identify novel inhibitors of inflammatory mediators (drug repurposing) but also to explore the influence of vitamin D and its existing analogues against PD through modulating inflammation. It might be possible to distinguish a few analogues as specific inhibitors or multitargeted inhibitors of the inflammatory pathway proteins.

## 2. Methods

For the current study we selected 10 analogues of vitamin-D (calcitriol, active form of vit-D), which are commercially available (see Table 1 for more details). On contrary, a few of the known inflammatory mediators were selected as targets form the literature.

**Table 1.**
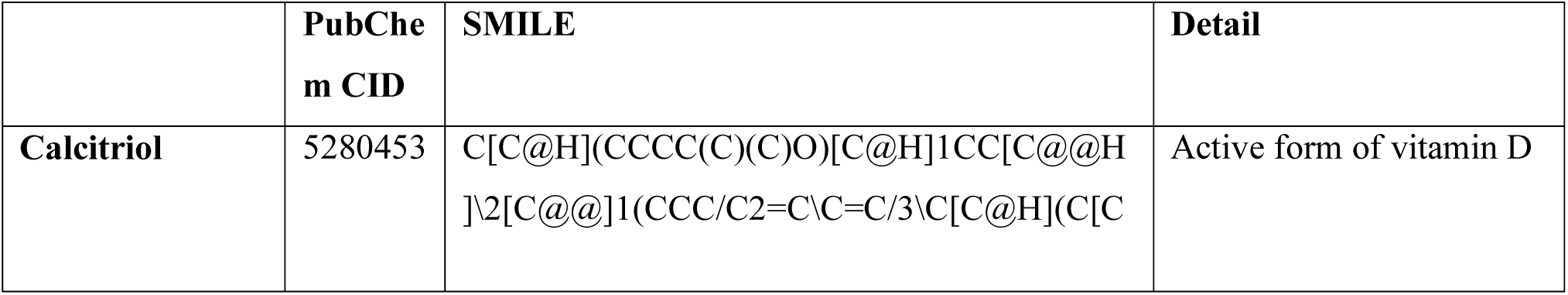
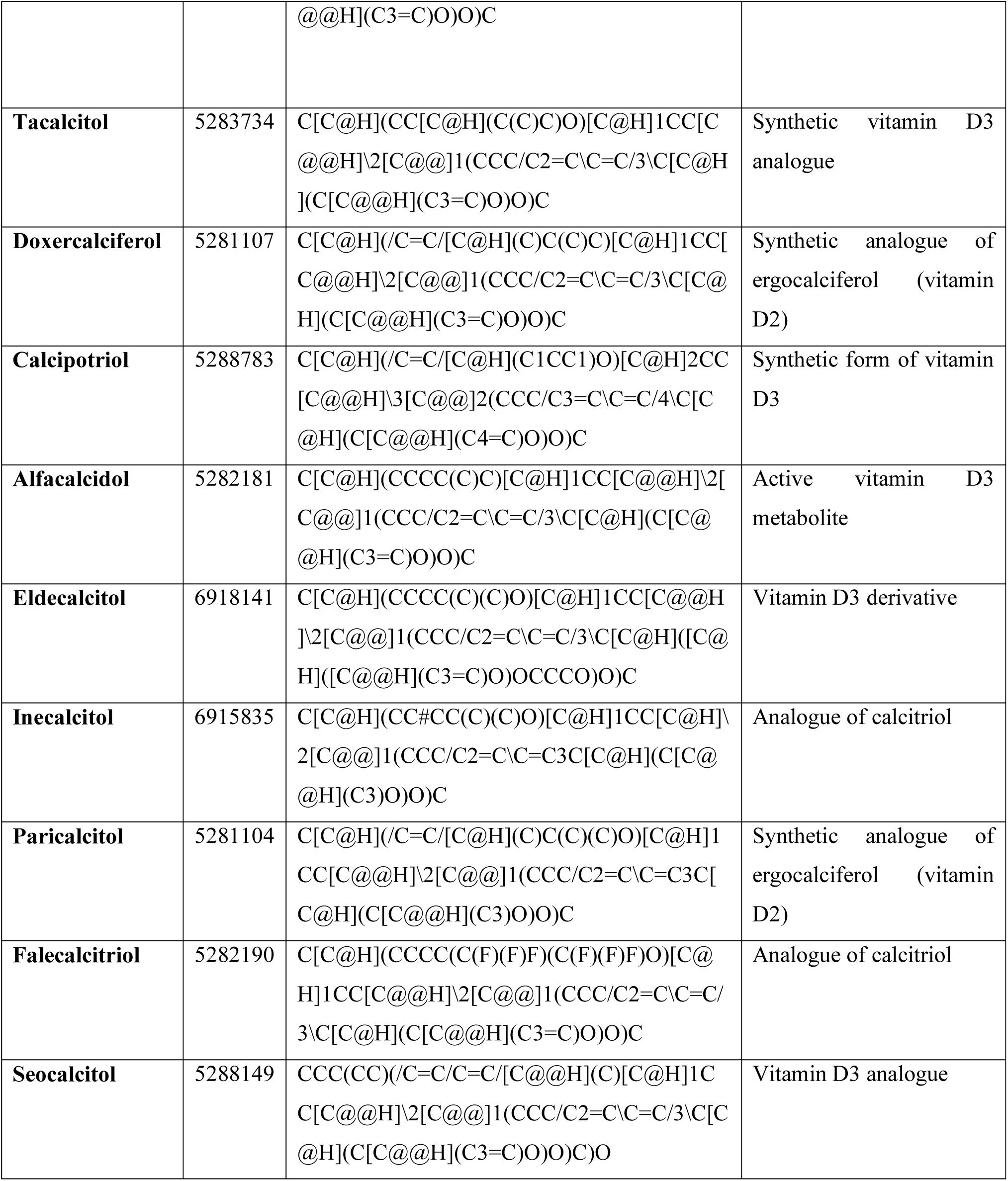
List of vitamin-D analogues used in this study as well as the chemical identifier and their nature.

### 2.1. Protein preparation procedure

The coordinates of the crystal structures of the target proteins were retrieved from the RCSB protein data bank (PDB) in pdb format. Missing atoms within the protein structure were repaired with the help of the Chimera (v1.18). These crystal structures were cleaned prior to the docking by removing the bound ligands, ions, and water molecules. The crystal structures of p38 MAPK, NF-κB, iNOS, nNOS, COX-2, TNFα, IL6, TNFα, IL1β, CRP were prepared for AutoDock Vina docking such as addition of polar hydrogens, charges, and assigning AD4 atom type. The preparation steps were carried out using AutoDock 4 (Morris et al., 2009).

### 2.2. Ligand preparation procedure

Three-dimensional structure of Vitamin D (calcitriol) and its selected analogues (Table 1), were retrieved in structure data file (SDF) format from the PubChem database. Energy minimisation of the selected ligands was performed with the help of the Avogadro (v1.2.0) software, and ligand files were saved as PDB format. Subsequently, PDB files were converted to pdbqt format using AutoDockTools (v1.5.7). For all the target proteins, we used either already bound ligand structure with the PDB file or standard drugs available (experimental or marketed).

### 2.3. Performing protein-ligand docking

Molecular docking was executed using AutoDock Vina 1.1.2. The grid box for docking was set in such a way that it can cover the whole protein. The compounds were screened based on Vina binding energy (Kcal/mol) and the interactions of ligands with target proteins were investigated with the help of Discovery Studio Visualizer.

### 2.4. Investigation of the blood-brain-barrier (BBB) permeability and neurotoxicity

Two different toxicity-related servers, Deep-PK (https://biosig.lab.uq.edu.au/deeppk/) and ProTox 3.0 (https://tox.charite.de/protox3/index.php?site=home) were utilised for analysis of the blood-brain-barrier permeability and neurotoxicity of the vitamin-D and its selected analogues.

## 3. Results

### 3.1. Vitamin-D and its analogues interacted with the p38 MAPK

When we screened for p38 MAPK modulators using the calcitriol and its analogues, we found calcipotriol and inecalcitol showed very good binding affinity compared to the selected reference compound, CHEMBL4574467 (-9.30 kcal/mol) (Table 2). Calcipotriol was bound with the p38 MAPK with a binding free energy of -9.20 kcal/mol (Table 2) and with two H-bonds, Thr107(4.13 Å), and Ile148 (5.47 Å) (Figure 1). On the other hand, inecalcitol showed lesser binding affinity (-9.70 kcal/mol) (Table 2) than the reference compound, while interacting with the kinase domain of the p38 MAPK, and forming one H-bond (Ile167, 4.78 Å) (Figure 1). Moreover, we didn’t find my any interaction between the reference compound and p38 MAPK which my influence the kinase activity of the enzyme. Whereas, inecalcitol and calcipotriol were able to interact with a few important residues within the kinase domain which regulate its activities. For example, inecalcitriol interacted with two ATP binding residues (Val31 and Tyr36), two residues (Ala52 and Lys54) which are very close to another ATP binding site of the enzyme (lys53), and proton acceptor site of the enzyme (Leu168) with three encompassing residues (Ile167, Asp169, and Phe170) (Figure 1). Similarly, calcipotriol also interacted with important residues for enzyme activity, including Tyr36 and Leu168 (Figure 1).

**Figure 1:**
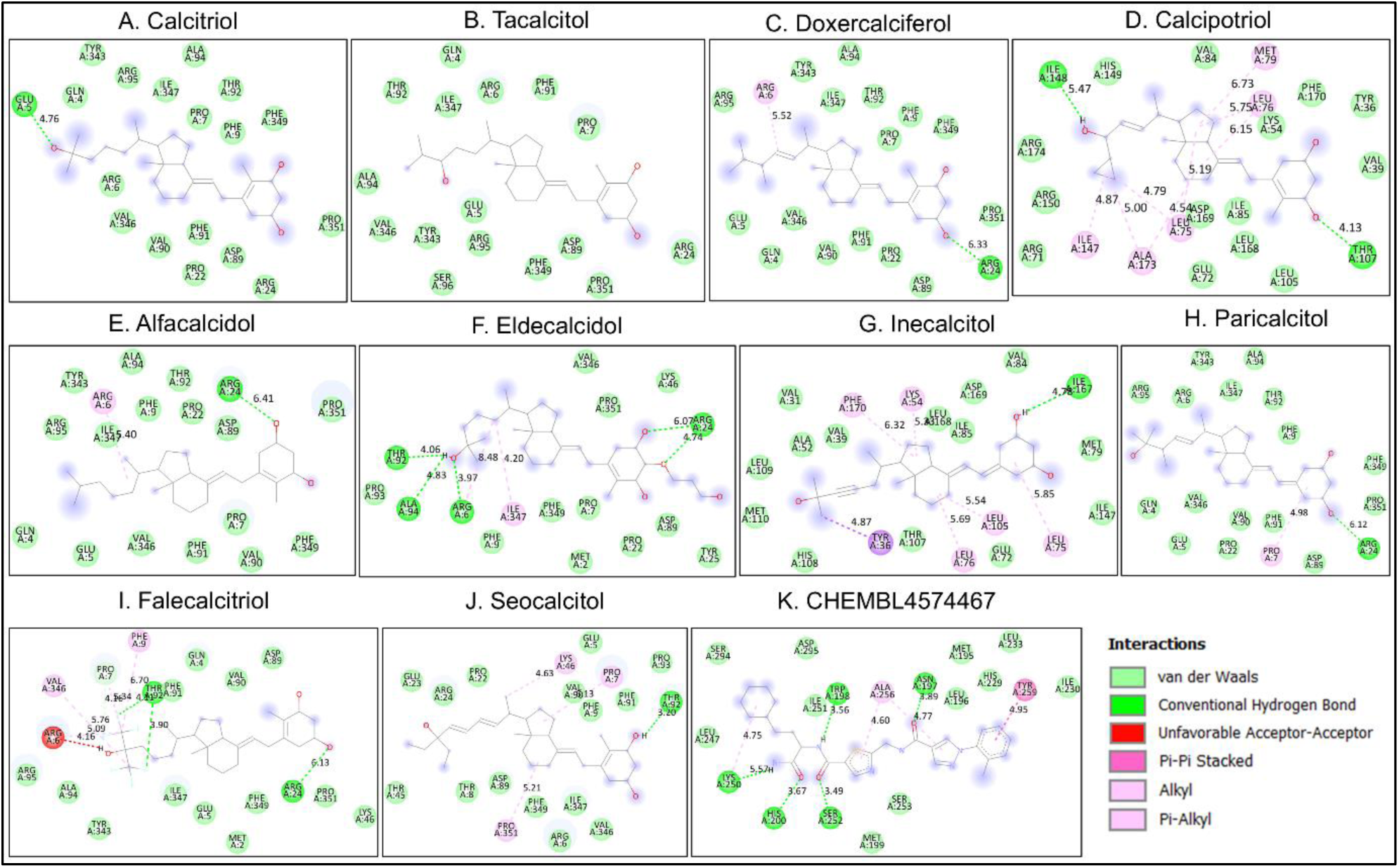
The Protein-ligand interactions were depicted from the molecular docking analysis (AutoDock Vina) between the p38 MAPK and calcitriol (with analogues). Interactions were visualised with the help of Discovery Studio which revealed bonded and non-bonded interactions, A) Calcitriol, B) Tacalcitol, C) Doxercalciferol, D) Calcipotriol, E) Alfacalcidol, F) Eldecalcidol, G) Inecalcitol, H) Paricalcitol, I) Falecalcitriol, J) Seoclcitol, and K) CHEMBL4574467.

**Table 2.**
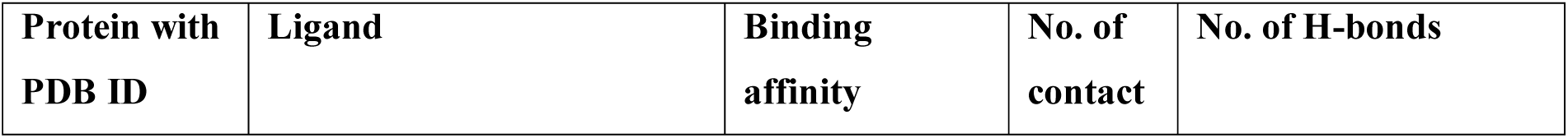
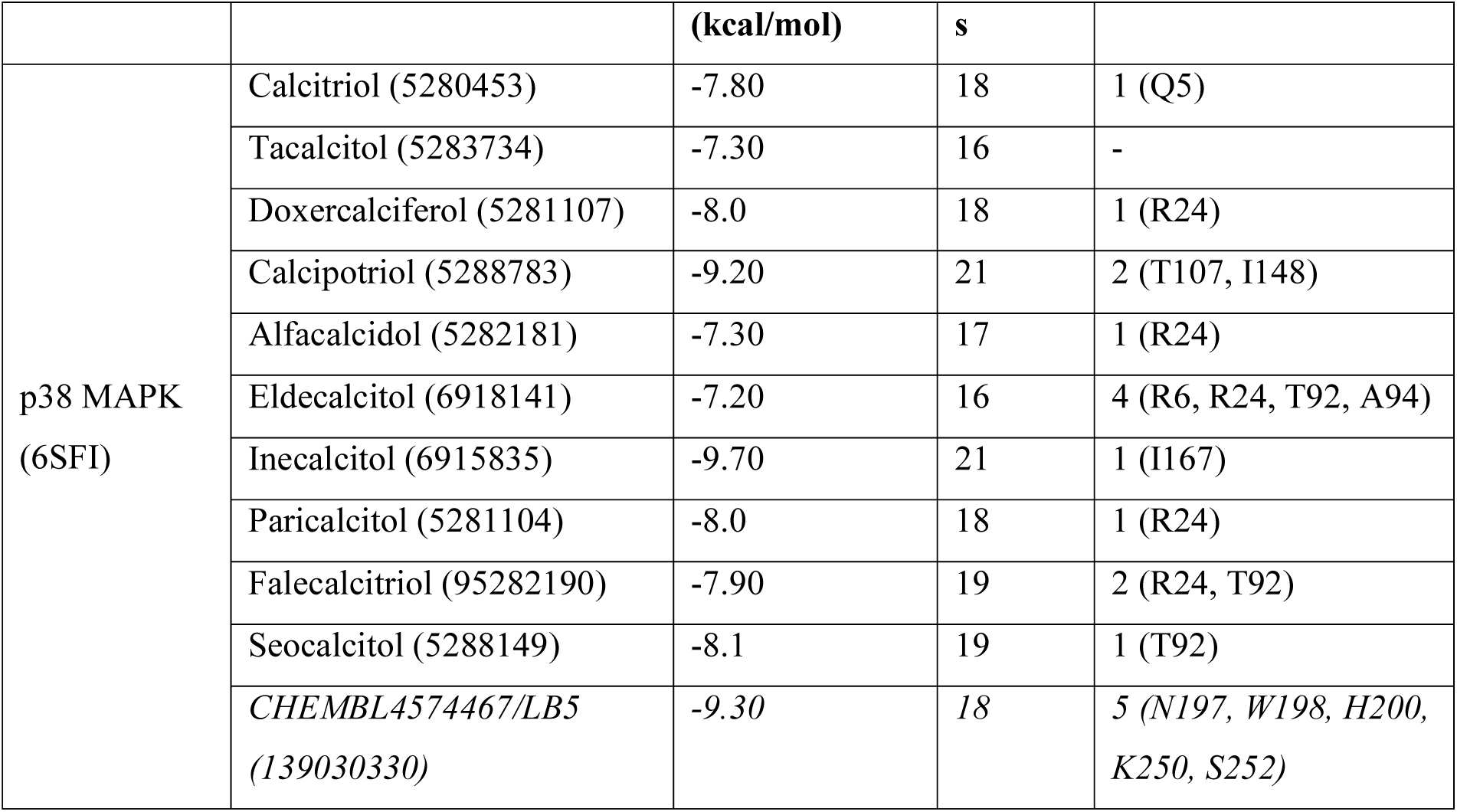
Details of the interaction between p38 MAPK and calcitriol, including its analogues. Reference compound used in molecular docking study highlighted in italic.

### 3.2. Vitamin-D and its analogues interacted with the Nuclear factor NF-kappa-B p105

Molecular docking analysis revealed that selected ligands were able to interact with several residues within the RHD domain (Rel homology domain) by forming H-bonds (occasionally) and non-covalent interactions (Figure 2). These compounds also showed modest binding affinity against the target protein which are very similar to the reference compound, ranging from -6.3 to -7.2 kcal/mol (Table 3). Seocalcitol showed the lowest binding free energy, thereby the highest binding affinity. However, alfacalcidol, eldecalcitol, inecalcitol, and falecalcitriol interacted with the target protein by engaging in maximum number of contacts through bonded as well as non-bonded interactions (Figure 2 and Table 3). Thereby these compounds might have some advantage over seocalcitol to modify the enzyme action.

**Figure 2:**
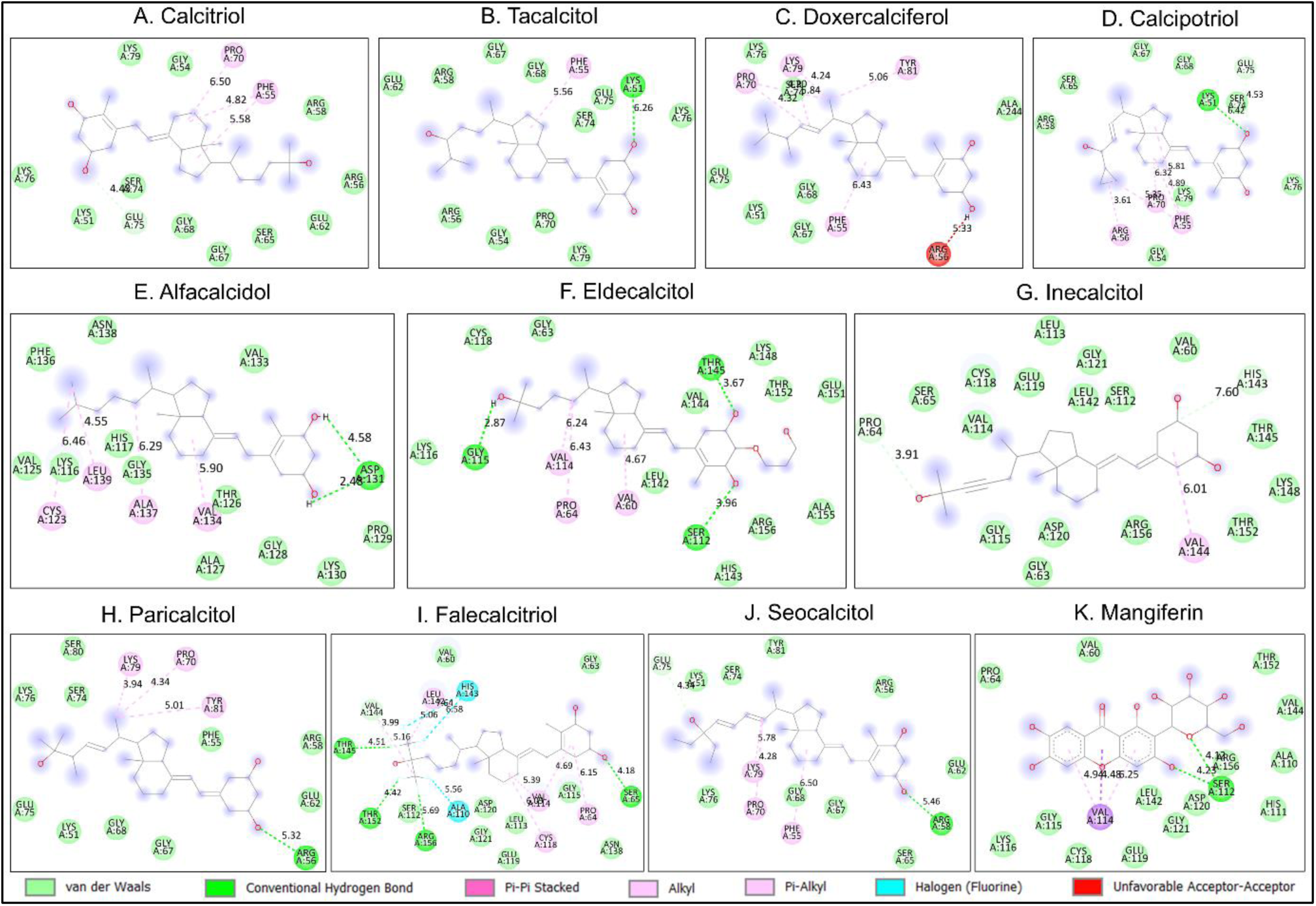
The Protein-ligand interactions were depicted from the molecular docking analysis (AutoDock Vina) between the nuclear factor NF-kappa-B p105 and calcitriol (with analogues). Interactions were visualised with the help of Discovery Studio which revealed bonded and non-bonded interactions, A) Calcitriol, B) Tacalcitol, C) Doxercalciferol, D) Calcipotriol, E) Alfacalcidol, F) Eldecalcidol, G) Inecalcitol, H) Paricalcitol, I) Falecalcitriol, J) Seoclcitol, and K) Mangiferin.

**Table 3.**
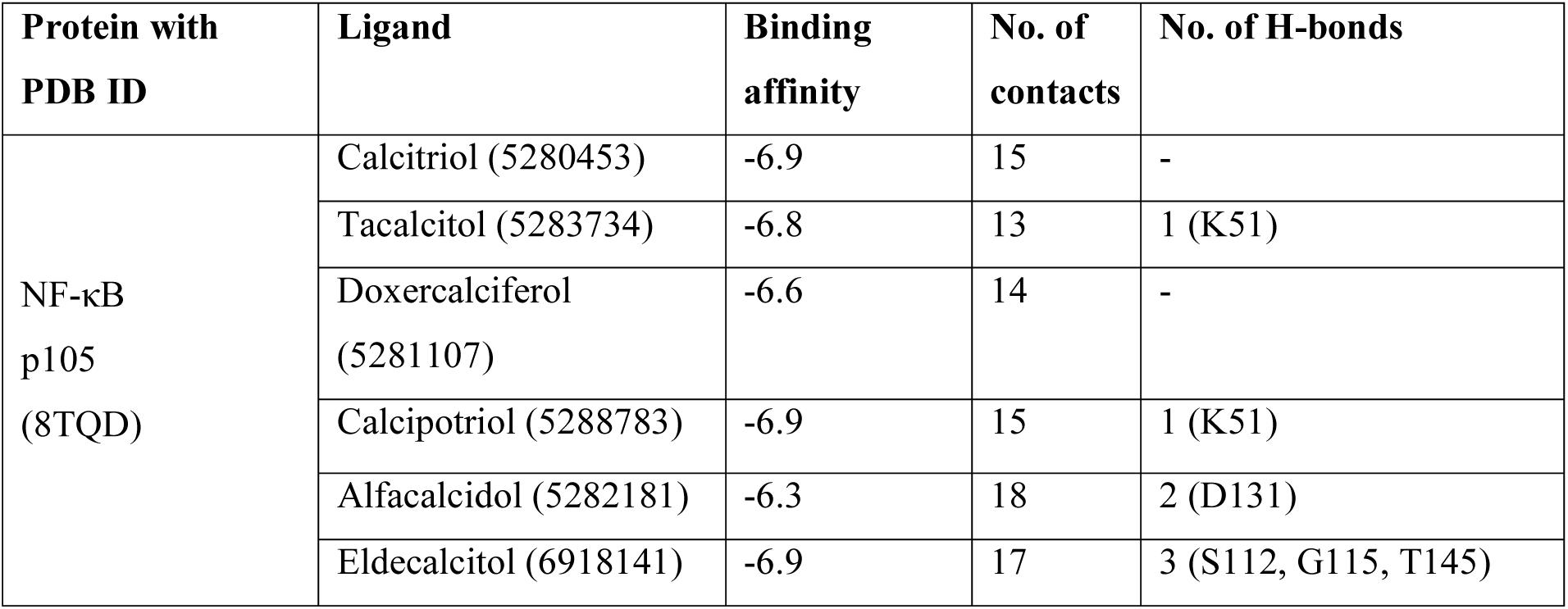
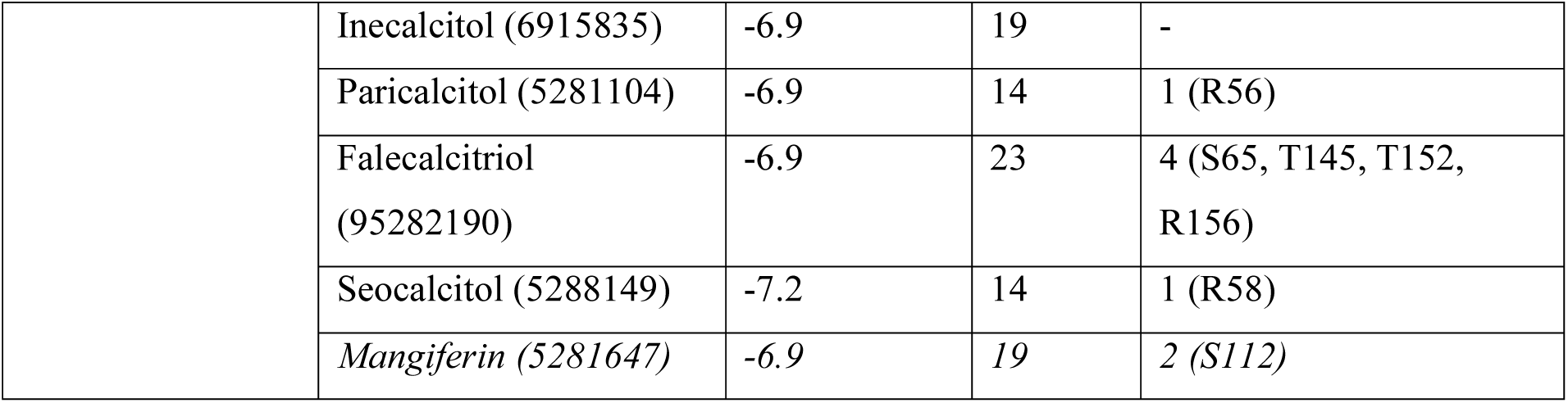
Details of the interaction between NF-kappa-B p105 and calcitriol, including its analogues. Reference compound used in molecular docking study highlighted in italic.

### 3.3. Vitamin-D and its analogues interacted with the iNOS and nNOS

All the tested ligands showed good binding affinity evidenced by negative binding free energy from the iNOS-ligand interactions, ranging from -8.1 to -9.5 kcal/mol (Table 4). Whereas, the reference compound showed only -6.9 kcal/mol binding free energy (Table 4). Thus, tested compounds might interact strongly with the target protein and have better chance to modulate the enzyme actions. The top three compounds based on binding free energy analysis were revealed to be paricalcitol (-9.5 kcal/mol), doxercalciferol (-9.4 kcal/mol), and calcipotriol (-9.2 kcal/mol) with a number of H-bonds, Alkyl, Pi-alkyl interaction, and Pi-Pi interaction (Figure 3). All of these three compounds were able to interact with residues (Trp113, Ala116, Pro117, Arg118, Cys119, Ile120, Trp382) (Figure 3) which are within very close proximity of the cofactor binding site of the enzyme. Calcitriol also showed very good binding affinity (-8.9 kcal/mol) towards the enzyme, iNOS (Table 4). Therefore, it is possible that these compounds might modify the enzyme activity of the iNOS.

**Figure 3:**
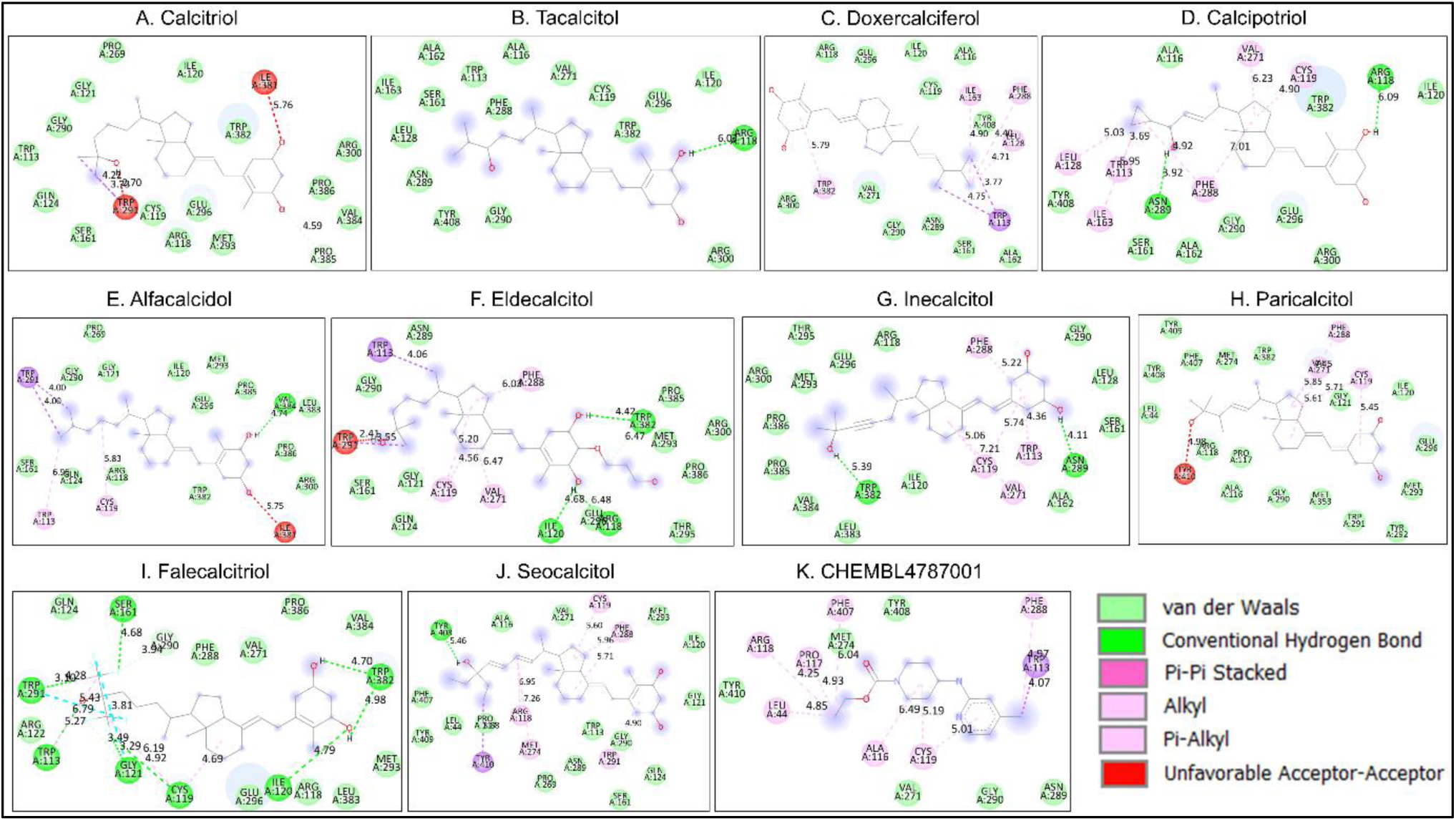
The Protein-ligand interactions were depicted from the molecular docking analysis (AutoDock Vina) between the iNOS and calcitriol (with analogues). Interactions were visualised with the help of Discovery Studio which revealed bonded and non-bonded interactions, A) Calcitriol, B) Tacalcitol, C) Doxercalciferol, D) Calcipotriol, E) Alfacalcidol, F) Eldecalcidol, G) Inecalcitol, H) Paricalcitol, I) Falecalcitriol, J) Seoclcitol, and K) CHEMBL4787001.

**Table 4.**
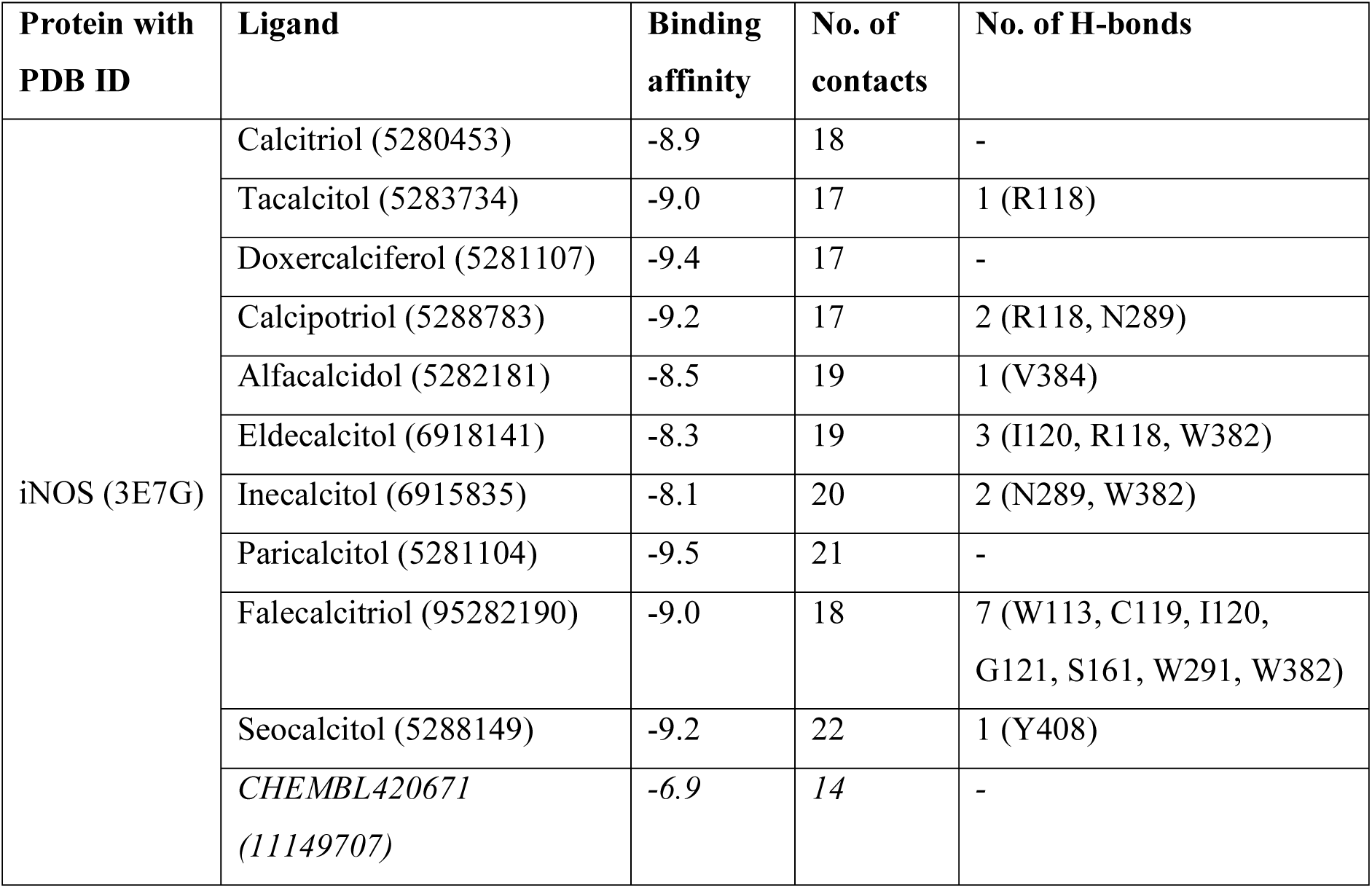
Details of the interaction between iNOS and calcitriol, including its analogues. Reference compound used in molecular docking study highlighted in italic.

| Protein with PDB ID | Ligand | Binding affinity | No. of contacts | No. of H-bonds |
| --- | --- | --- | --- | --- |
| iNOS (3E7G) | Calcitriol (5280453) | -8.9 | 18 | - |
|  | Tacalcitol (5283734) | -9.0 | 17 | 1 (R118) |
|  | Doxercalciferol (5281107) | -9.4 | 17 | - |
|  | Calcipotriol (5288783) | -9.2 | 17 | 2 (R118, N289) |
|  | Alfacalcidol (5282181) | -8.5 | 19 | 1 (V384) |
|  | Eldecalcitol (6918141) | -8.3 | 19 | 3 (I120, R118, W382) |
|  | Inecalcitol (6915835) | -8.1 | 20 | 2 (N289, W382) |
|  | Paricalcitol (5281104) | -9.5 | 21 | - |
|  | Falecalcitriol (95282190) | -9.0 | 18 | 7 (W113, C119, I120, G121, S161, W291, W382) |
|  | Seocalcitol (5288149) | -9.2 | 22 | 1 (Y408) |
|  | *CHEMBL420671*<br>(11149707) | -6.9 | 14 | - |

Coming to the nNOS, all of the ligands showed decent binding affinity (binding energy range, -8.9 to - 10.3 kcal/mol) towards the target protein (Table 5) and also formed a number of H-bonds and other non-bonded interactions (Figure 4).

**Figure 4:**
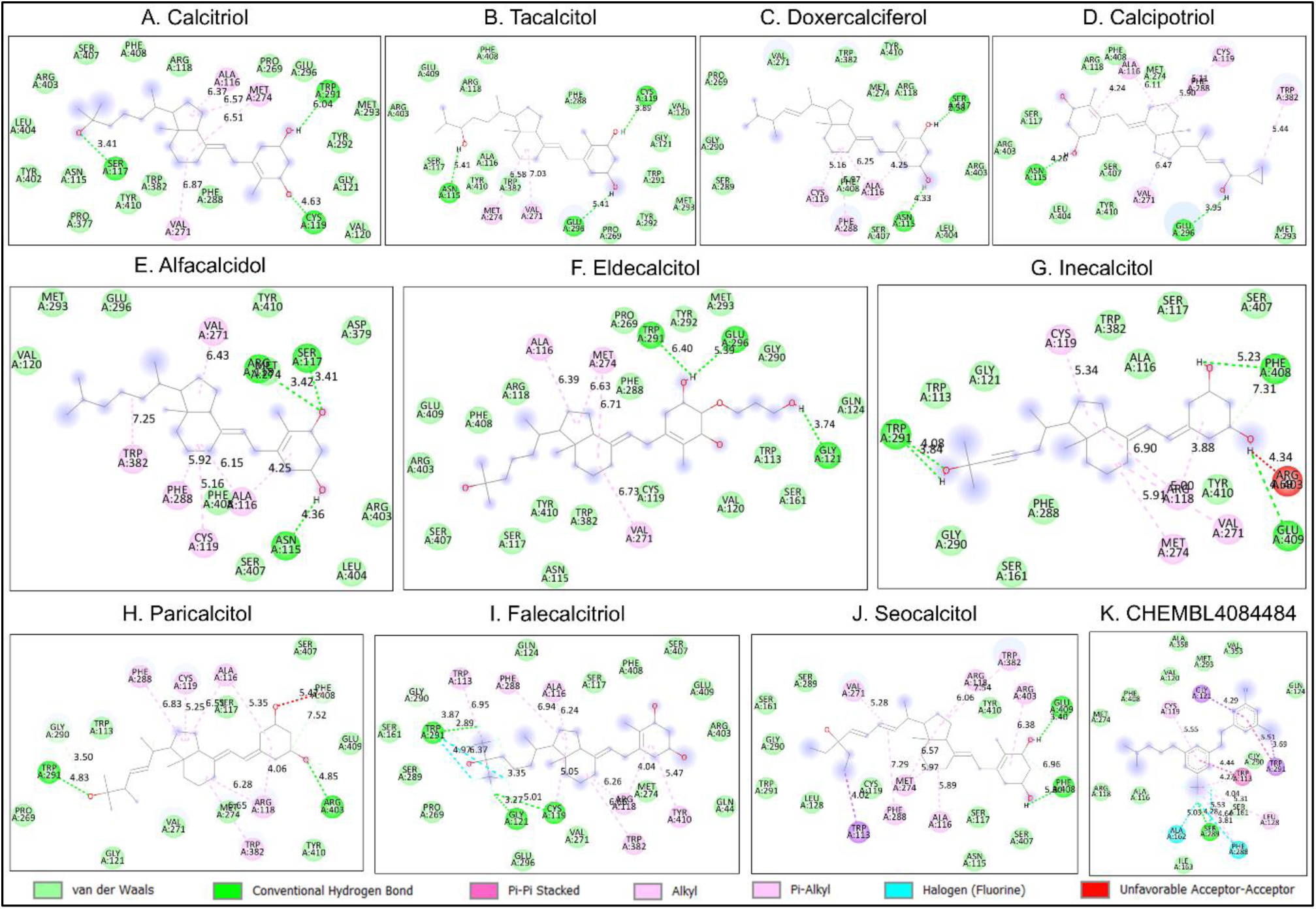
The Protein-ligand interactions were depicted from the molecular docking analysis (AutoDock Vina) between the nNOS and calcitriol (with analogues). Interactions were visualised with the help of Discovery Studio which revealed bonded and non-bonded interactions, A) Calcitriol, B) Tacalcitol, C) Doxercalciferol, D) Calcipotriol, E) Alfacalcidol, F) Eldecalcidol, G) Inecalcitol, H) Paricalcitol, I) Falecalcitriol, J) Seoclcitol, and K) CHEMBL4084484.

**Table 5.** Details of the interaction between nNOS and calcitriol, including its analogues. Reference compound used in molecular docking study highlighted in italic.

| Protein | Ligand | Binding affinity | No. of contacts | No. of H-bonds |
| --- | --- | --- | --- | --- |
| nNOS (6AV2) | Calcitriol (5280453) | -9.6 | 24 | 3 (S117, C119, W291) |
|  | Tacalcitol (5283734) | -9.4 | 20 | 3 (N115, C119, E296) |
|  | Doxercalciferol (5281107) | -9.8 | 18 | 2 (N115, S117) |
|  | Calcipotriol (5288783) | -9.2 | 17 | 2 (N115, E296) |
|  | Alfalcidol (5282181) | -9.0 | 19 | 3 (N115, S117, R118) |
|  | Eldecalcitol (6918141) | -8.9 | 26 | 3 (G121, W291, E296) |
|  | Inecalcitol (6915835) | -9.2 | 21 | 4 (W291, F408, E409) |
|  | Paricalcitol (5281104) | -9.8 | 21 | 2 (W291, R403) |
|  | Falecalcitriol (95282190) | -9.7 | 28 | 3 (C119, G121, W291) |
|  | Seocalcitol (5288149) | -10.3 | 21 | 2 (F408, E409) |
|  | *CHEMBL4084484*<br>(134686795) | -9.5 | 24 | 1 (S289) |

### 3.4. Vitamin-D and its analogues interacted with COX-2

Calcitriol and all of its analogues showed either better or similar binding affinity when docked against the COX-2 active site (Table 6). Calcipotriol, alfacalcidol, and seocalcitol revealed as the top three modulators of the enzyme, with binding free energy ranging from -8.8 to -9.5 kcal/mol compared to the aspirin (reference drug, -7.3 kcal/mol) (Table 6) and calcipotriol was involved in highest number of contact points (residues). More interestingly we found calcipotriol and alfacalcidol interacted with the residues forming the second pocket (Val523, Tyr355, Leu352, Ser353, Pe518) inside the COX-2 active site and also the residue Phe518 which is part of the molecular gate that extends to the second drug binding pocket exclusively present in COX-2 (Figure 5). Contrastingly, we found calcitriol, tacalcitol, and doxercalciferol interacted with Arg120, which is the starting residue of the arachidonate binding site of the enzyme. (Figure 5). Overall, these compounds have the potential to modulate the COX-2 enzyme action either as a dual COX-1 and COX-2 inhibitor or specific COX-2 inhibitor.

**Figure 5:**
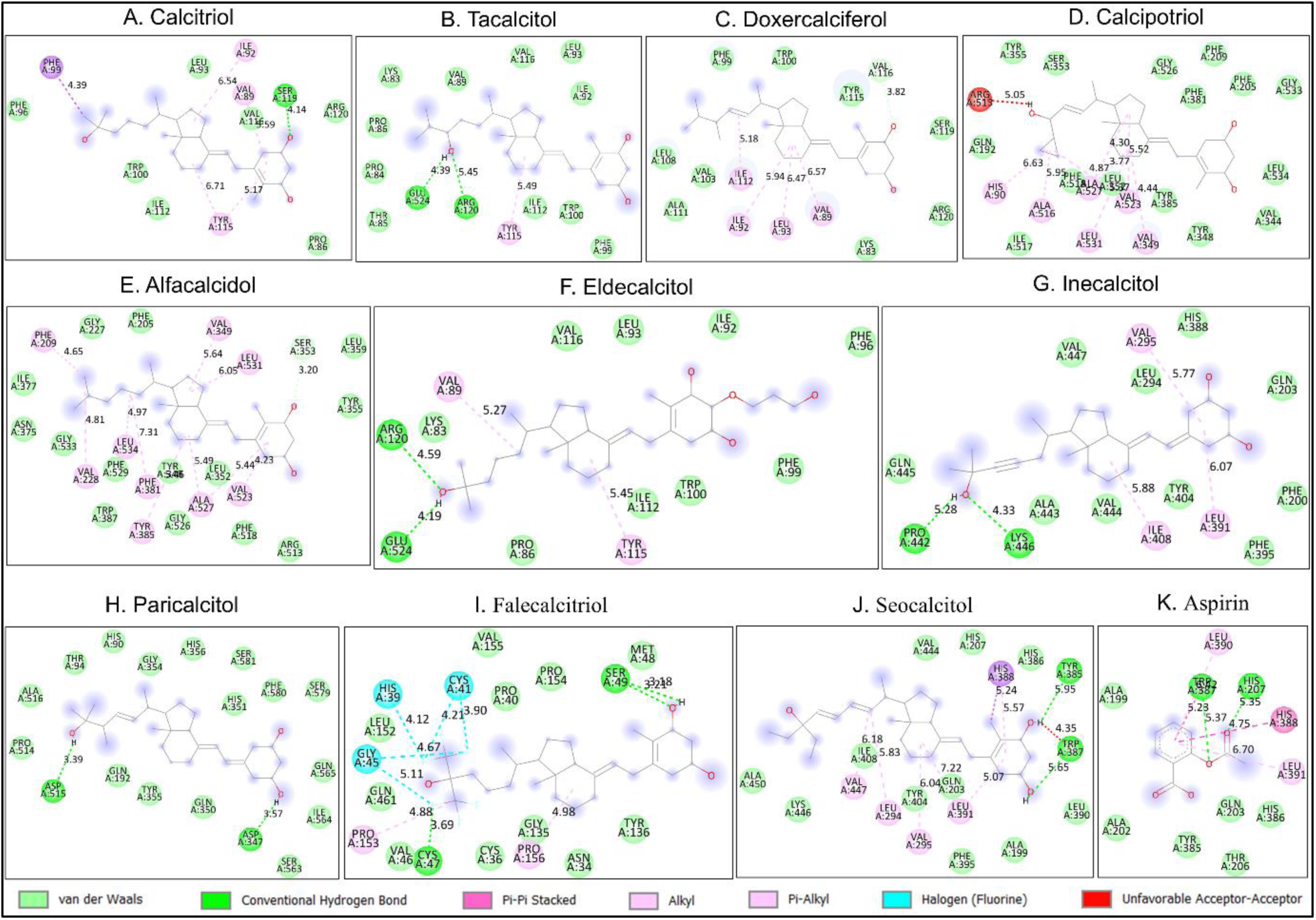
The Protein-ligand interactions were depicted from the molecular docking analysis (AutoDock Vina) between the COX-2 and calcitriol (with analogues). Interactions were visualised with the help of Discovery Studio which revealed bonded and non-bonded interactions, A) Calcitriol, B) Tacalcitol, C) Doxercalciferol, D) Calcipotriol, E) Alfacalcidol, F) Eldecalcidol, G) Inecalcitol, H) Paricalcitol, I) Falecalcitriol, J) Seoclcitol, and K) CHEMBL4084484.

**Table 6.**
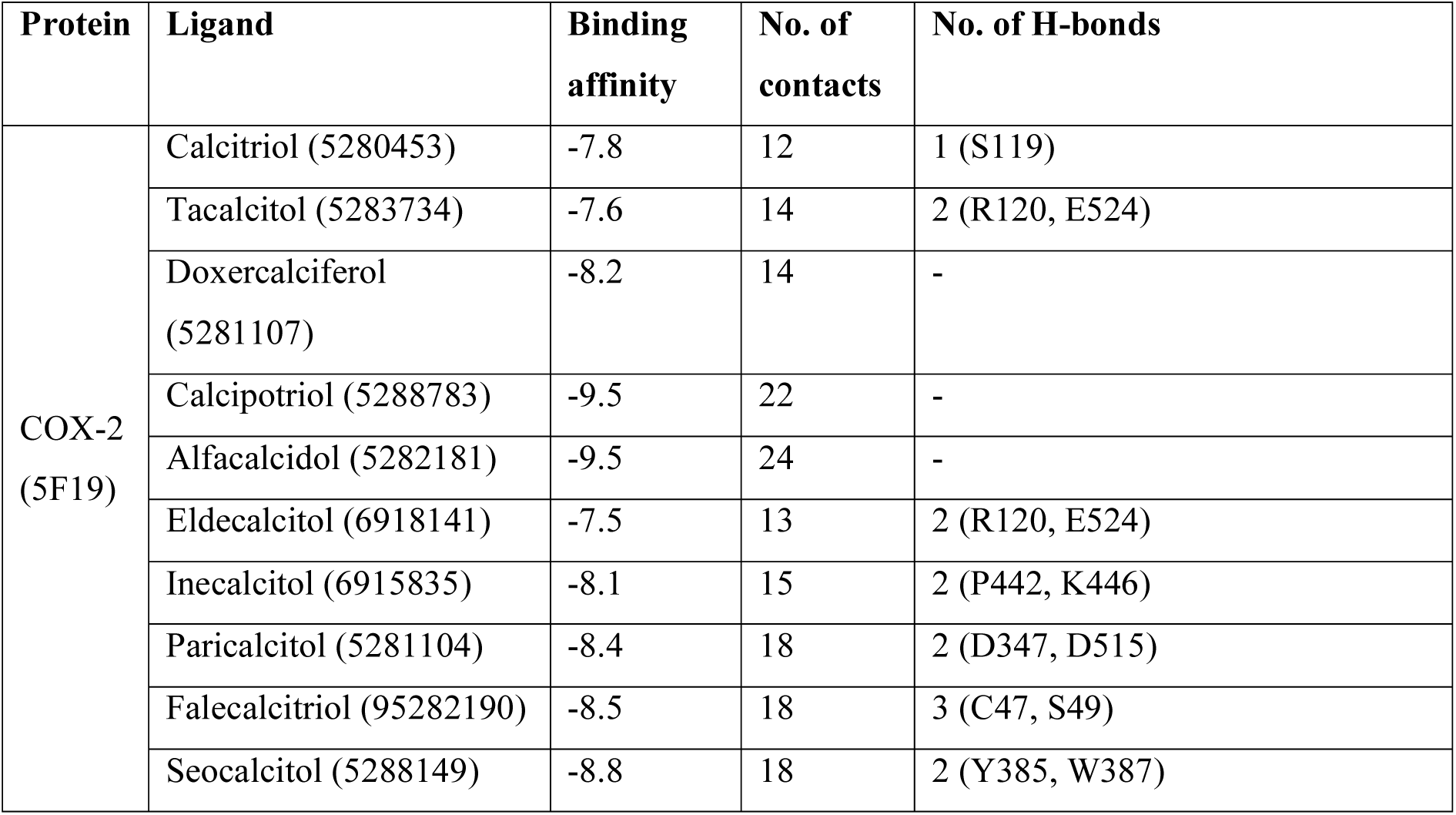
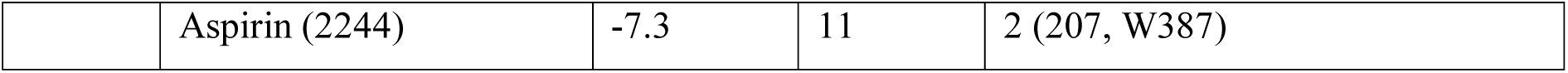
Details of the interaction between COX-2 and calcitriol, including its analogues. Reference compound used in molecular docking study highlighted in italic.

### 3.5. Vitamin-D and its analogues interacted with TNF-α

Tested compounds showed modest binding affinity towards the TNF-α, and only three compounds calcipotriol, paricalcitol, and seocalcitol exhibited better or same binding affinity (-7.0 to -9.2 kcal/mol) compared to the reference compound (CHEMBL4787001, -7.0 kcal/mol) (Table 7). However, none of these compounds were able to interact with the important residues which are linked to biological activities of TNF-α. Thereby, these compounds might interact with separate binding sites within the active site of the TNF-α to modulate its function. However, calcitriol, tacalcitol, doxercalciferol, and falecalcitriol were able to interact with residues like Gln105, Thr108, and Ala112 of TNF-α, which are very important for the enzyme activity. Not only that, a few compounds which are located very close proximity to these residues (Pro103, Cys104, Arg106, Glu107, Pro109, Glu113, Ala114, Lys115, Pro116, Trp117) and might have better possibility to modify enzyme action (Figure 6).

**Figure 6:**
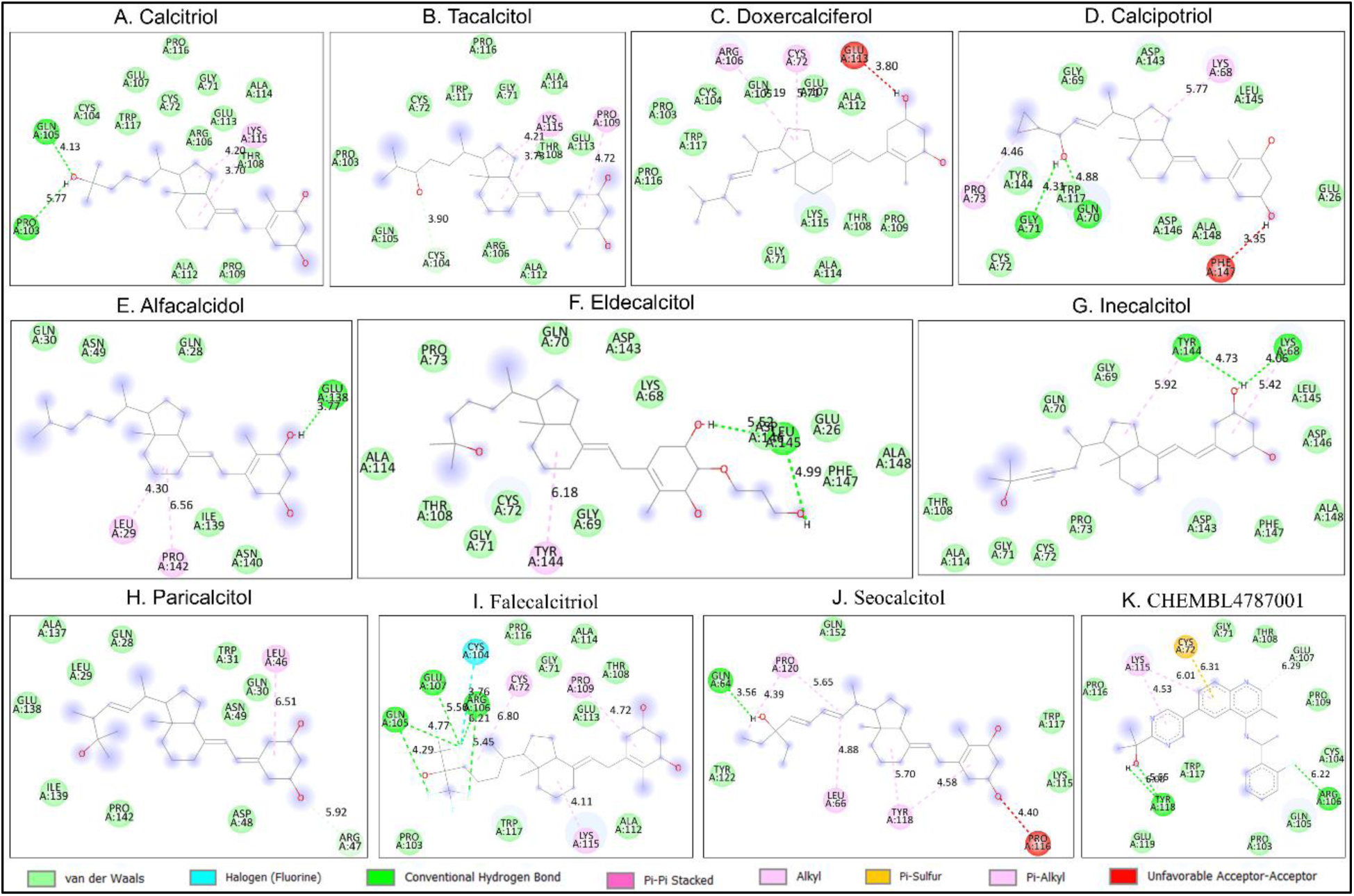
The Protein-ligand interactions were depicted from the molecular docking analysis (AutoDock Vina) between the TNF-α and calcitriol (with analogues). Interactions were visualised with the help of Discovery Studio which revealed bonded and non-bonded interactions, A) Calcitriol, B) Tacalcitol, C) Doxercalciferol, D) Calcipotriol, E) Alfacalcidol, F) Eldecalcidol, G) Inecalcitol, H) Paricalcitol, I) Falecalcitriol, J) Seoclcitol, and K) CHEMBL4084484.

**Table 7.** Details of the interaction between TNF-α and calcitriol, including its analogues. Reference compound used in molecular docking study highlighted in italic.

| Protein | Ligand | Binding affinity | No. of contacts | No. of H-bonds |
| --- | --- | --- | --- | --- |
| TNFα (7JRA) | Calcitriol (5280453) | -6.20 | 15 | 2 (P103, Q105) |
|  | Tacalcitol (5283734) | -6.00 | 14 | - |
|  | Doxercalciferol (5281107) | -6.5 | 15 | - |
|  | Calcipotriol (5288783) | -9.20 | 14 | 2 (Q70, G71) |
|  | Alfalcacidol (5282181) | -5.70 | 8 | 1 (E138) |
|  | Eldecalcitol (6918141) | -6.60 | 15 | 2 (L145) |
|  | Inecalcitol (6915835) | -6.60 | 14 | 2 (K68, Y144) |
|  | Paricalcitol (5281104) | -7.50 | 12 | - |
|  | Falecalcitriol (95282190) | -6.60 | 15 | 3 (Q105, R106, E107) |
|  | Seocalcitol (5288149) | -7.00 | 9 | 1 (Q64) |
|  | CHEMBL4787001 (126531356) | -7.0 | 14 | 2 (R106, Y118) |

### 3.6. Vitamin-D and its analogues interacted with IL-6

Calcitriol itself and its analogues showed better binding affinity than the reference compound, berberine (Table 8). Paricalcitol came out as the best ligand to modify the IL-6 action in terms of binding affinity, whereas, calcipotriol and inecalcitol were involved in maximum number of contacts, same as the paricalcitol (14) (Table 8). However, we could not find any interactions between the tested ligands and the target protein, involving key residues (Tyr32, Phe74, Gln75, Val104, Leu178, Arg179, Gln175, and Gln183) (Figure 7) which are previously reported as essential for the protein function. Therefore, it might be a possibility that these compound act on different site of the IL-6 to modify its action, provided these compounds actually modify the protein function (subject to in vitro assay).

**Figure 7:**
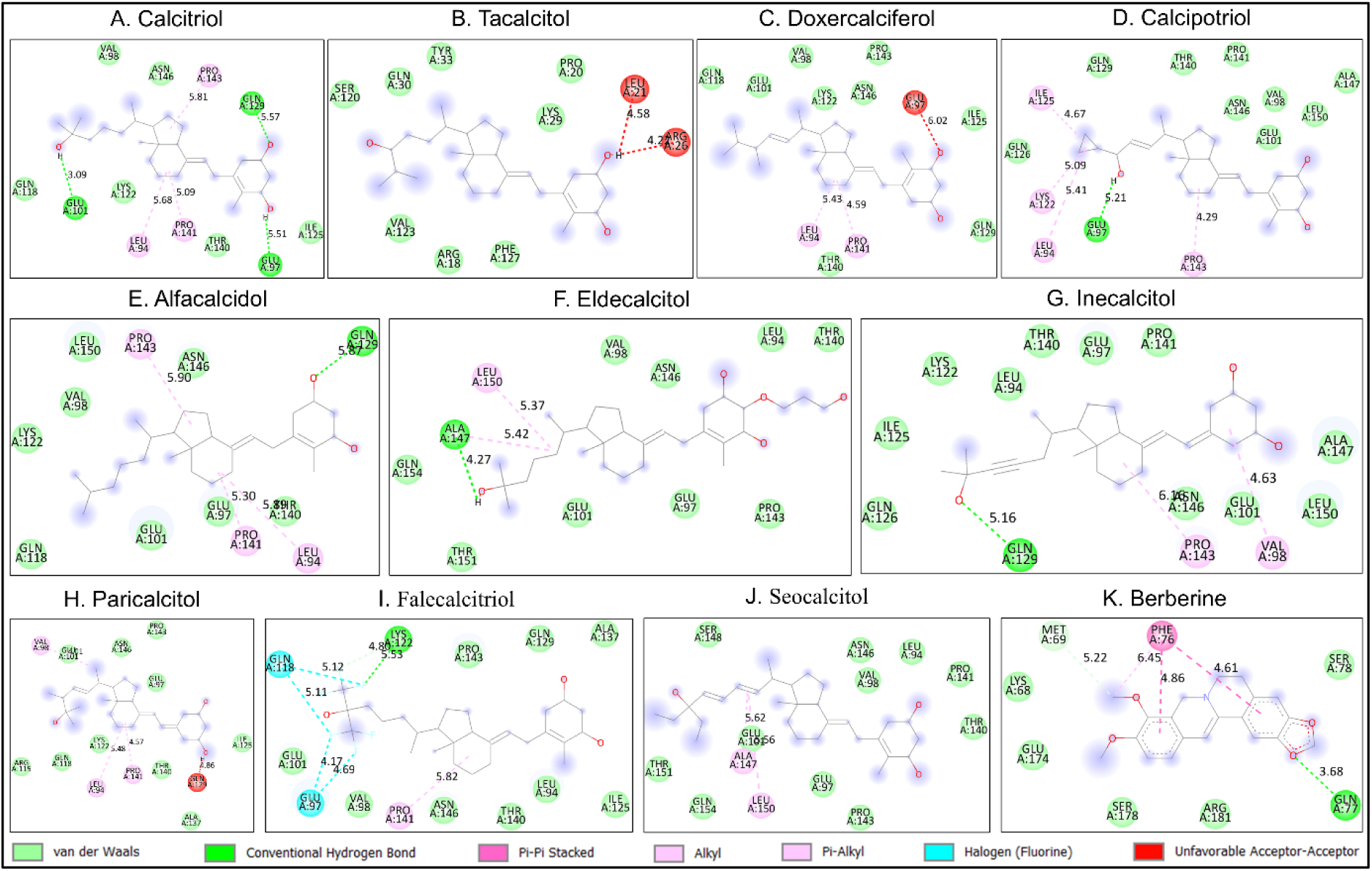
The Protein-ligand interactions were depicted from the molecular docking analysis (AutoDock Vina) between the IL-6 and calcitriol (with analogues). Interactions were visualised with the help of Discovery Studio which revealed bonded and non-bonded interactions, A) Calcitriol, B) Tacalcitol, C) Doxercalciferol, D) Calcipotriol, E) Alfacalcidol, F) Eldecalcidol, G) Inecalcitol, H) Paricalcitol, I) Falecalcitriol, J) Seoclcitol, and K) CHEMBL4084484.

**Table 8.** Details of the interaction between IL-6 and calcitriol, including its analogues. Reference compound used in molecular docking study highlighted in italic.

| Protein | Ligand | Binding affinity | No. of contacts | No. of H-bonds |
| --- | --- | --- | --- | --- |
| IL6<br>(4NI7) | Calcitriol (5280453) | -6.9 | 12 | 3 (E97, E101, Q129) |
|  | Tacalcitol (5283734) | -6.2 | 10 | - |
|  | Doxercalciferol<br>(5281107) | -7.2 | 12 | - |
|  | Calcipotriol (5288783) | -6.8 | 14 | 1 (E97) |
|  | Alfacalcidol (5282181) | -6.7 | 12 | 1 (Q129) |
|  | Eldecalcitol (6918141) | -6.3 | 11 | 1 (A147) |
|  | Inecalcitol (6915835) | -6.5 | 14 | 1 (Q129) |
|  | Paricalcitol (5281104) | -7.4 | 14 | - |
|  | Falecalcitriol<br>(95282190) | -6.7 | 13 | 1 (K122) |
|  | Seocalcitol (5288149) | -7.0 | 13 | - |
|  | Berberine (2353) | -6.6 | 8 | 1 (Q77) |

### 3.7. Vitamin-D and its analogues interacted with IL-1β

When we checked for interaction between protein and ligand in the case of IL-1β, we found calcitriol and its analogues showed less binding affinity towards the target protein, but a few compounds showed good binding affinity (Table 8) which involved many bonded and non-bonded interactions (Figure 8). Doxercalciferol, calcipotriol, alfacalcidol, inecalcitol, paricalcitol, and falecalcitriol showed binding free energy less than or equal to -7.0 kcal/mol, which indicated possible interference of the protein function in presence of these compounds (Table 9). IL-1β contains two active sites, known as site A and site B. We found the reference compound T9C and calcitriol were engaged in contacts with residues (Val3, Phe46, Ile56, Lys92, Lys93, and Lys94 of the site B (Il-1β) (Figure 8). In contrast, paricalcitol interacted with site A residues (Ser125, Gln126, Ala127, Met130, and Pro131) and inecalcitol interacted with Met20, Ser21, Gly22, and Gln38 (Figure 8). Additionally, a few other compounds, including tacalcitol, doxercalciferol, calcipotriol, and alfacidol were interacted with both site A and site B of the protein.

**Figure 8:**
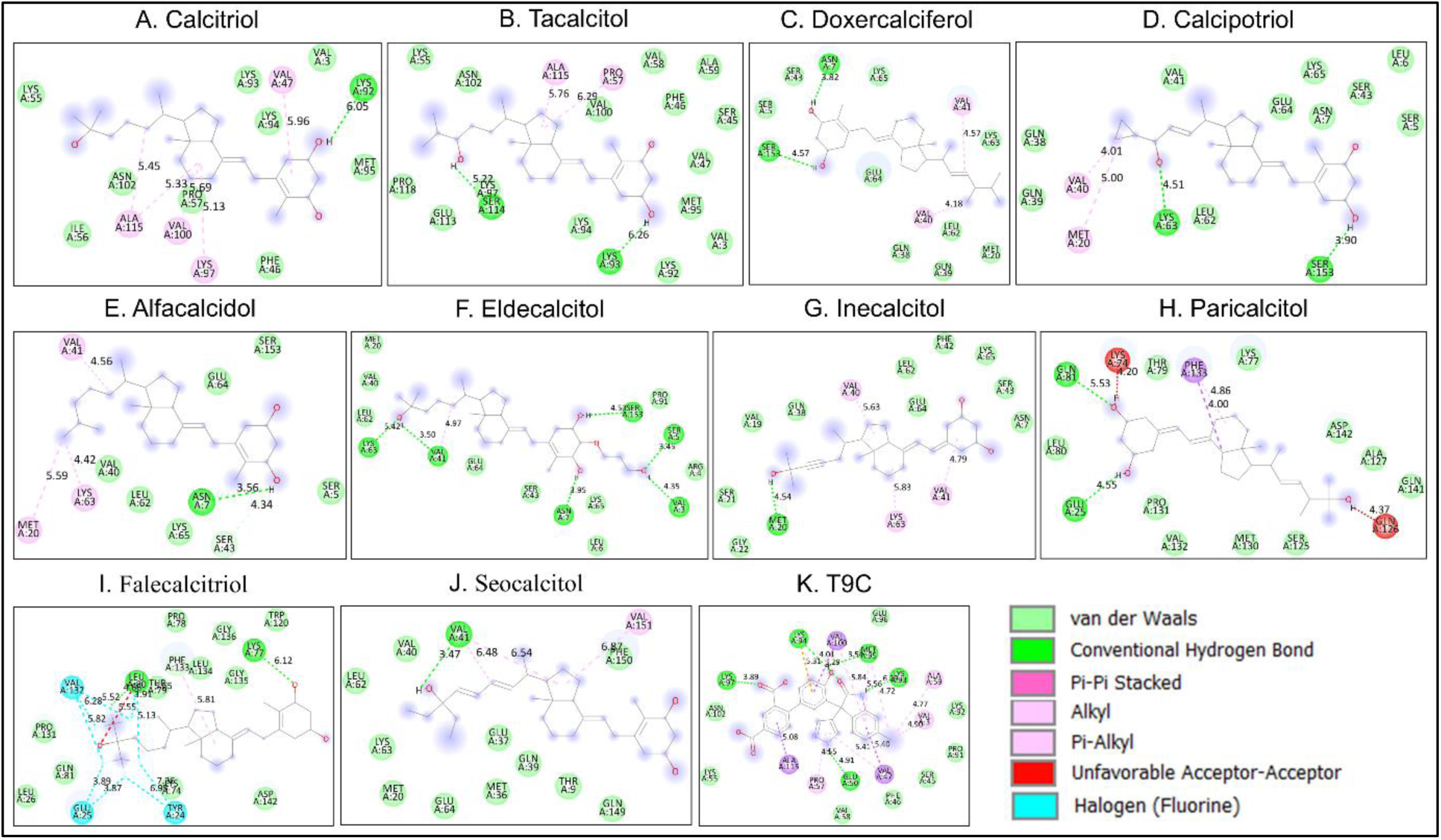
The Protein-ligand interactions were depicted from the molecular docking analysis (AutoDock Vina) between the IL-1β and calcitriol (with analogues). Interactions were visualised with the help of Discovery Studio which revealed bonded and non-bonded interactions, A) Calcitriol, B) Tacalcitol, C) Doxercalciferol, D) Calcipotriol, E) Alfacalcidol, F) Eldecalcidol, G) Inecalcitol, H) Paricalcitol, I) Falecalcitriol, J) Seoclcitol, and K) CHEMBL4084484.

**Table 9.**
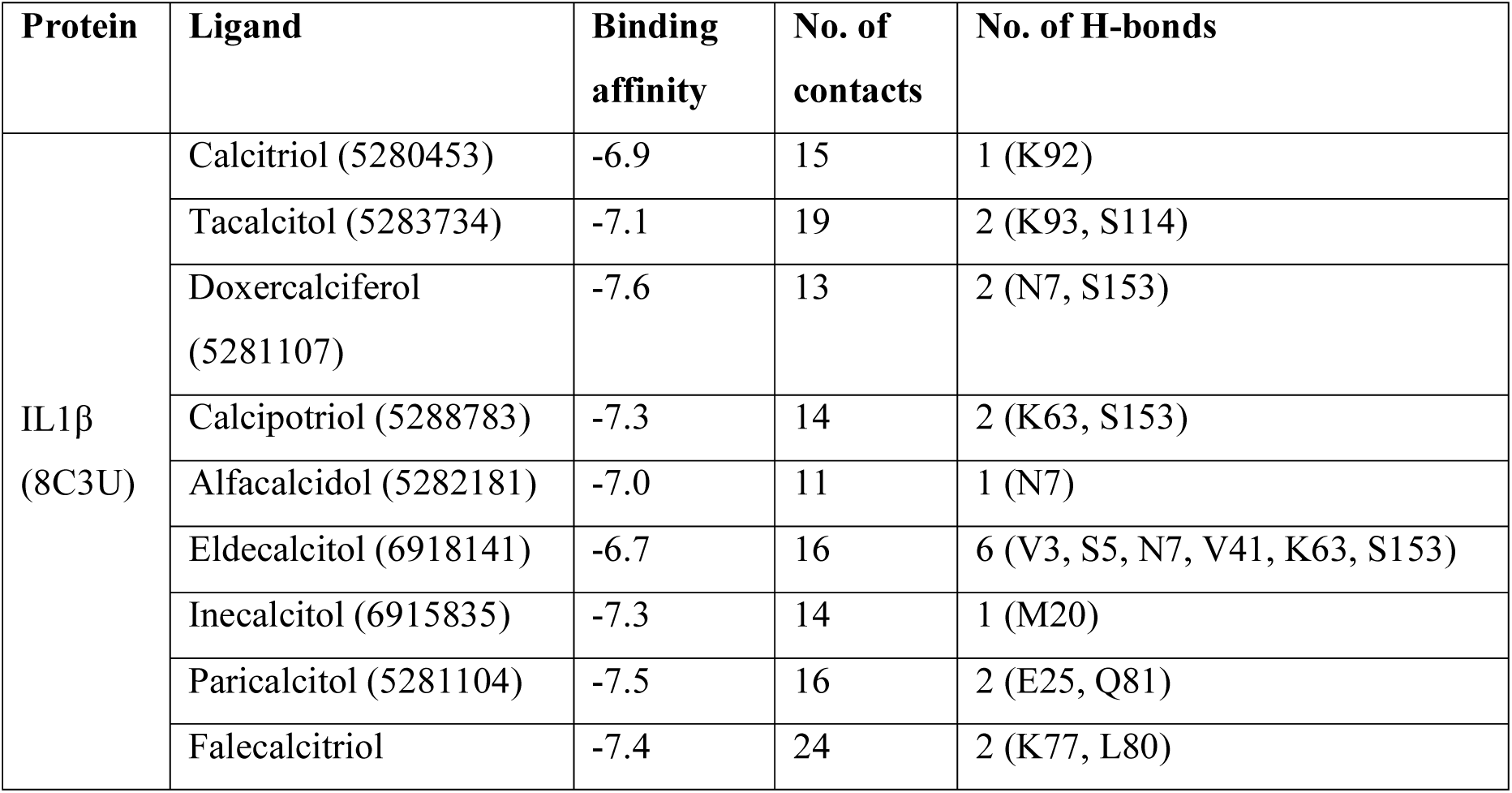
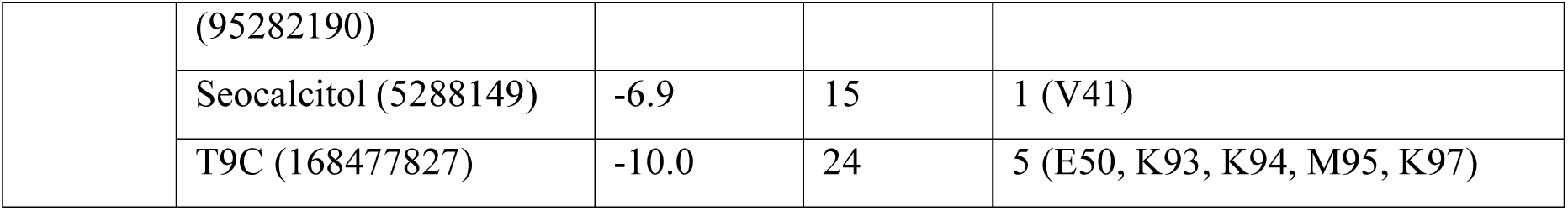
Details of the interaction between IL-1β and calcitriol, including its analogues. Reference compound used in molecular docking study highlighted in italic.

### 3.8. Vitamin-D and its analogues interacted with CRP

In the case of CRP, we found a few showed much better affinity than the reference compound (Table 10). For example, falecalcitriol, paricalcitol, doxercalciferol, inecalcitol, and calcipotriol showed binding affinity less than -8.0 kcal/mol, compared to reference compound (-4.9 kcal/mol) (Table 9). Falecalcitriol was involved in maximum number of contact points as well as formed 5 H-bonds (Figure 7). Other protein-ligand interactions also involved H-bonds and non-covalent interactions (Figure 7). However, we didn’t find any interactions between protein and ligand which involves calcium binding residues (Asp78, Asn79, Glu156, Gln157, Asp158, and Gln168). Therefore, these compounds bind with the CRP chain without affecting the calcium binding and their mode of action might be something else.

**Table 10.** Details of the interaction between CRP and calcitriol, including its analogues. Reference compound used in molecular docking study highlighted in italic.

| Protein | Ligand | Binding affinity | No. of contacts | No. of H-bonds |
| --- | --- | --- | --- | --- |
| CRP<br>(1B09) | Calcitriol (5280453) | -7.9 | 19 | 1 (T41) |
|  | Tacalcitol (5283734) | -7.9 | 17 | - |
|  | Doxercalciferol<br>(5281107) | -8.4 | 18 | - |
|  | Calcipotriol (5288783) | -8.2 | 18 | 1 (T41) |
|  | Alfacalcidol (5282181) | -7.6 | 16 | 1 (P115) |
|  | Eldecalcitol (6918141) | -7.2 | 19 | 3 (E85, T90) |
|  | Inecalcitol (6915835) | -8.3 | 21 | 1 (T41) |
|  | Paricalcitol (5281104) | -8.4 | 17 | 1 (S44) |
|  | Falecalcitriol<br>(95282190) | -8.7 | 22 | 5 (S44, Y49, Y73, A92) |
|  | Seocalcitol (5288149) | -7.9 | 17 | 1 (A92) |
|  | Phosphocholine (1014) | -4.9 | 16 | 5 (T41, S44, Y49, T90, A92) |

**Figure 9:**
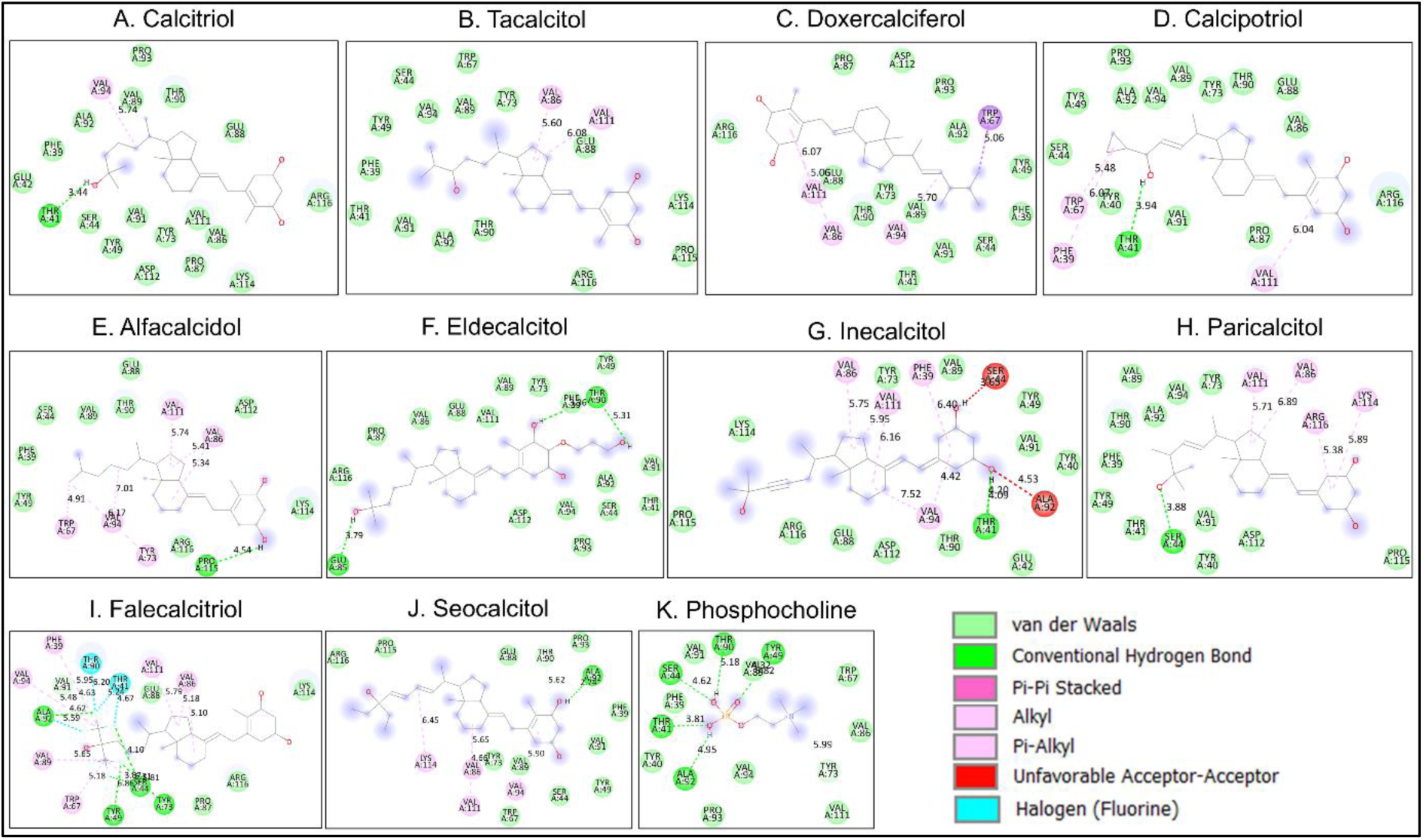
The Protein-ligand interactions were depicted from the molecular docking analysis (AutoDock Vina) between the IL-1β and calcitriol (with analogues). Interactions were visualised with the help of Discovery Studio which revealed bonded and non-bonded interactions, A) Calcitriol, B) Tacalcitol, C) Doxercalciferol, D) Calcipotriol, E) Alfacalcidol, F) Eldecalcidol, G) Inecalcitol, H) Paricalcitol, I) Falecalcitriol, J) Seoclcitol, and K) CHEMBL4084484

### 3.9. Most of the ligands are non-neurotoxic and can pass BBB

When we checked for the BBB permeability status for the vitamin-D and its analogues we found most of the compounds were predicted to be BBB permeable by both the servers. However, in the case of tacalcitol, eldecalcitol, and inecalcitol there were disparities between the prediction of two different servers (Table 11). Therefore, another server was used to get more clarity, admetSAR 3.0 (https://lmmd.ecust.edu.cn/admetsar3/index.php). For tacalcitol and inecalcitol the BBB permeability was found to be active, 60% and 61.4% respectively. Whereas, eldecalcitol was found to be non-permeable (0%). Therefore, except eldecalcitol, two other compounds tacalcitol and inecalcitol might cross BBB as suggested by two different servers. On top that, none of these compounds were associated with neurotoxicity (Table 11).

**Table 11.** Neurotoxicity status and Blood-brain-barrier (BBB) permeability of calcitriol and its analogues.

|  | Deep-pk |  | BBB-barrier (ProTox 3.0) |  | Neurotoxicity (ProTox 3.0) |  |
| --- | --- | --- | --- | --- | --- | --- |
|  | Blood-brain barrier | Predictive confidence | Prediction | Probability | Prediction | Probability |
| **Calcitriol** | Penetrable<br>(Medium Confidence) | 0.795 | Active | 0.59 | Inactive | 0.94 |
| **Tacalcitol** | Non-Penetrable<br>(Medium Confidence) | 0.265 | Active | 0.59 | Inactive | 0.94 |
| **Doxercalciferol** | Penetrable<br>(High Confidence) | 0.958 | Active | 0.67 | Inactive | 0.67 |
| **Calcipotriol** | Penetrable<br>(Medium Confidence) | 0.694 | Active | 0.67 | Inactive | 0.93 |
| **Alfacalcidol** | Penetrable<br>(High Confidence) | 0.918 | Active | 0.66 | Inactive | 0.69 |
| **Eldecalcitol** | Non-Penetrable<br>(High Confidence) | 0.036 | Active | 0.71 | Inactive | 0.97 |
| **Inecalcitol** | Non-Penetrable<br>(High Confidence) | 0.01 | Active | 0.64 | Inactive | 0.85 |
| **Paricalcitol** | Penetrable<br>(Medium Confidence) | 0.796 | Active | 0.66 | Inactive | 0.89 |
| **Falecalcitriol** | Penetrable<br>(High Confidence) | 1.0 | Active | 0.81 | Inactive | 0.92 |
| **Seocalcitol** | Penetrable<br>(Low<br>Confidence) | 0.526 | Active | 0.68 | Inactive | 0.84 |

## 4. Discussion

Chronic inflammation is zenith of reasons behind many diseases, including neurodegenerative diseases. There are several players in the inflammatory pathway which have been previously reported in the pathophysiology of Parkinson’s disease. NF-κB and p38 MAPK are two indispensable regulators of the inflammation. Here, we found our two best ligands, inecalcitol and calcipotriol interacted strongly with the p38 MAPK, and specifically involved in ATP binding (Val31, Tyr36, and Lys53) and proton acceptor site (Leu168) (Figure 1). Apart from that, these compounds were also interacted with several other important residues, important for protein function (Figure 1). Calcitriol itself also showed modest binding affinity. Calcitriol was previously reported to ameliorate the p38 MAPK signalling in mice hepatorenal toxicity (Morsy *et al*, 2021) and shown anti-atherosclerotic properties in LPS-induced inflammation in HUVEC human cells via inhibition of NF-κB and p38 MAPK (Talmor *et al*, 2008). We also found from the docking study, calcitriol and its analogues were able to interact with the RHD domain of the p50 subunit of the NF-κB protein. Thus, it is quite possible that a few ligands, including calcitriol itself may modify the NF-κB function to combat inflammation (Figure 2). Calcitriol and some of its analogues also possess anti-cancer properties thanks to their abilities to the MAPK function (Segovia-Mendoza *et al*, 2015). From the protein-ligand interaction, a few analogues were revealed to be potential modulator of the p38 MAPK and NF-κB and which might contribute to their anti-inflammatory activities.

Anti-inflammatory activities of calcitriol might be through COX and NOS enzymes. Calcitriol is linked to COX-1 and COX-2 inhibition to suppress inflammatory response (Suo *et al*, 2021; Wang, *et al*, 2014). Further, calcitriol has previously shown to influence the NOS activity (Andrukhova *et al*, 2014; Willems *et al*, 2012). In the current study, we found possible molecular interactions between these inflammatory pathway enzymes and calcitriol and its analogues. Not only that, a few of the compounds (paricalcitol, doxercalciferol, and calcipotriol) alongside calcitriol were involved in strong interactions (Table 4) with the iNOS protein and interacted with cofactor binding sites (Figure 3). Thereby, these compounds might be strong candidates to modify enzyme actions by interfering with the cofactor binding. Apart from that, calcitriol and its analogues showed strong binding affinity towards the nNOS enzyme.

Calcipotriol, alfacalcidol, and seocalcitol revealed as the top three modulators of the COX-2 (Table 6). Calcipotriol and alfacalcidol interacted with the second pocket (Val523, Tyr355, Leu352, Ser353, Pe518) of the COX-2 active site, and also the residue Phe518 which acts as the molecular gate of the second drug binding pocket in COX-2 (Figure 5). Contrastingly, a few compounds (calcitriol, tacalcitol, and doxercalciferol) interacted with Arg120, (arachidonate binding site residue) (Figure 5). Thus, these compounds potentially modulate the COX-2 enzyme action to repress inflammation.

A delicate balance between pro and anti-inflammatory cytokines drives the inflammation and any imbalance leads to issue in the resolution phase of the inflammation and may result in chronic inflammation which is substantially linked to PD (Dzamko *et al*, 2023). Pro-inflammatory cytokines play a crucial role (TNFα, IL-6, IL-1β) in PD, and exact role of these cytokines are still complicated (Dzamko *et al*, 2023). Vitamin D is related to the modulation of neuroinflammation and here we found calcitriol and its analogues were involved in strong interaction with pro-inflammatory cytokines. Although, calcitriol or its analogues were not able to interact with the TNF-α active site, but calcitriol, tacalcitol, doxercalciferol, and falecalcitriol may modify the protein activity through Gln105, Thr108, and Ala112, which are related to the proper function of the TNF-α (Figure 6). Previously, serum 25(OH)D status was shown to be negatively correlated to TNF-α concentrations in healthy women and may contribute to Vitamin-D’s role as an anti-inflammatory agent (Peterson and Heffernan, 2008). In the case of IL-6, none of the ligands interacted with the well-characterised residues known to be important for the IL-6 function (Figure 7 and Table 7). However, those compounds interacted strongly with the IL-6, compared to the reference compound. Additionally, previous works reported anti-inflammatory action of Vitamin D might be trough suppressing IL-6. From the protein-ligand interaction analysis, it was revealed that the strong interaction between calcitriol and its analogues with the IL-6 might be through different site rather than the known drug binding pocket. Calcitriol and analogues were also involved in interaction with either site A or site B and sometimes both sites of IL-1β active sites (Figure 8). Even though the binding affinity of these compounds were lesser than the reference compound (Table 9), but their interaction with crucial residues (Val3, Gln126, Phe46, Ile56, Lys92, Lys93, Lys94, Ser125, Ala127, Met130, and Pro131), within the active sites of the enzyme might be enough to modulate the IL-1β function. In line with our findings, previously Vitamin-D administration was reported to negatively regulate IL-1β, upon LPS (lipopolysaccharide) exposure (Ge *et al*, 2019). Not only that, calcipotriol and alfacalcidol also suppressed IL-1β production in different experimental set up (in vitro and in vivo) (Grimm *et al*, 2017). On the other hand, tacalcitol inhibited IL-6 and IL-8 expression previously in human fibroblast cells (Rostkowska-Nadolska *et al*, 2010). Figure 10 is a comprehensive representation of the possible interactions of inflammatory mediators and the various VD analogues studied. Overall, these compounds along with the Vitamin-D might be crucial to find out novel functions of Vitamin-D in neuroinflammation and allied diseases like PD. It would also be possible to find out novel drug activity of these compounds against PD.

**Figure 10:**
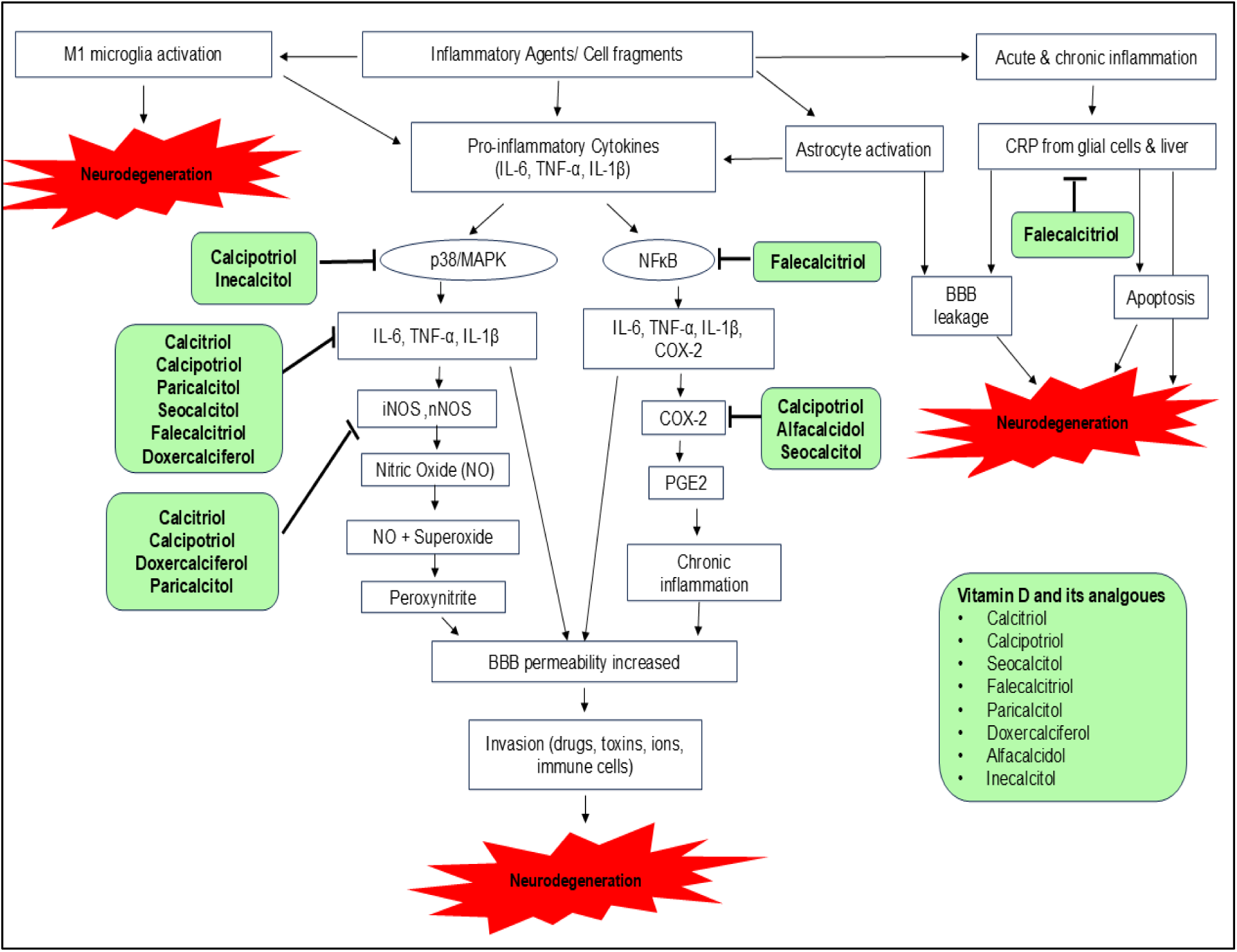
Possible interactions between calcitriol and its analogues with inflammatory mediators.

This *in-silico* approach offers a cost-effective and effectual method to ascertain novel functions of Vitamin D and its analogues in modulating neuroinflammation, giving a deeper insight of their role in various neurodegenerative diseases. Additionally, it provides potential for drug discovery by simulating interactions and predicting the biological activity of these compounds, opening possibilities for developing new therapies for conditions like Parkinson’s disease. While many possible interactions might be found by the predictive algorithms, not all of them might be clinically significant. Certain interactions could be overstated or misunderstood in the absence of biological context.

## 5. Conclusion

This study provides a comprehensive representation of the potential interactions between inflammatory mediators and VD and its analogues, offering acumen on their role in neuroinflammation and related diseases, such as Parkinson’s disease. This study suggests that biologically active VD (calcitriol) along with VD analogues, may hold promise in uncovering novel functions of vitamin D in regulating neuroinflammatory processes. Moreover, this approach highlights the potential of naturally occurring compounds like calcitriol or similar compounds which might have therapeutic effect against neuroinflammatory aspects of PD, paving the way for advanced experimental research and drug development and discovery targeting neurodegenerative disorders.

## Acknowledgment

Authors would like to acknowledge Dipsikha Khamrai for her efforts in helping with molecular docking protocols.

## Data Availability

Data and material will be made available to the corresponding author of the paper for review or query upon request.

## Author Contribution

R.P. and B.R. conceived and planned the manuscript. R.P. did the literature review and molecular docking. Manuscript writing, table and figures were prepared by R.P. The manuscript was critically checked and edited by N.D. B.R. critically checked, edited and validate the manuscript. All the authors read and approved the final manuscript.

## List of Abbreviations

BBB: Blood brain barrier
COX-1: Cyclooxygenase 1
COX-2: Cyclooxygenase 2
CRP: C reactive protein
IL-1β: Interleukin 1 beta
IL-6: Interleukin 6
IL-8: Interleukin 8
iNOS: inducible Nitric Oxide Synthase
MAPK: Mitogen Activated Protein Kinase
NF-κB: Nuclear Factor -kappa-B
nNOS: neuronal nitric oxide synthase
NO: nitric oxide
NOS: nitric oxide synthase
PD: Parkinson’s disease
PGE2: Prostaglandin E2
TLR-4: Toll-like receptor 4
TNFα: Tumour necrosis factor alpha
VD: vitamin D

